# Split intein vectors permit oversize FLIM-FRET biosensor neuronal expression

**DOI:** 10.64898/2026.06.02.725428

**Authors:** Jeong Oen Lee, Henry H. Puhl, Abigail Holder, Aurora Sheridan, Caleb Darden, Muhammad Shah, William Dunne, Tuan Nguyen, Kirk Hines, Shana Augustin, Youngchan Kim, Steven S. Vogel, David M. Lovinger

## Abstract

Adeno-associated virus (AAV) packaging limits constrain the design, performance, and in vivo application of genetically encoded biosensors. We developed a split intein-mediated reconstitution strategy enabling modular delivery and reassembly of oversized fluorescence lifetime-based biosensors. Using this platform, we engineered an oversized cAMP sensor compatible with one-photon fluorescence lifetime measurements, enabling monitoring of intracellular signaling dynamics in distinct neuronal subtypes in freely behaving mice.

## MAIN

Genetically encoded fluorescent biosensors enable cell-type-specific measurement of intracellular signaling and neurotransmitter dynamics in vivo. Recent advances in fluorescence intensity-based sensors [1,2] have come to dominate in vivo neuroscience applications, owing to their high signal-to-noise ratios and compatibility with adeno-associated virus (AAV) delivery. However, optimizing these sensors requires extensive mutagenesis and screening, making it difficult to establish generalizable design strategies [3,4].

In contrast, Förster resonance energy transfer (FRET)-based biosensors [5] have been widely used in cell biology and biophysical studies because of their modular architecture and predictable photophysical properties. They also provide absolute fluorescence lifetime readouts, independent of optical systems and much less sensitive to expression levels, making them particularly attractive for long-term in vivo neural recording.

Despite these strengths and the availability of many FRET biosensor templates [6], AAV packaging constraints (∼4.7 kb) prevent their direct use in vivo. For example, FRET-based cAMP sensors exceed the size limit for AAV delivery and must be re-engineered into intensity-based formats to achieve compatibility [7,8]. This size constraint represents a general barrier in biosensor design, and to date, no broadly applicable strategies exist for adapting oversized FRET biosensors to AAV-compatible formats.

Here, we present a protein engineering strategy to overcome AAV packaging constraints by leveraging split intein-mediated protein splicing. Split inteins are evolutionarily conserved protein elements [9] that catalyze precise post-translational ligation of separately expressed protein fragments, a strategy that has recently been validated in AAV-based gene therapy [10,11]. We applied this approach to FRET biosensors that are otherwise difficult to deliver via AAV due to large sensing domains, such as the cAMP-binding domain of EPAC (exchange protein directly activated by cAMP). The designed FRET biosensor construct, comprising fluorescent protein components and EPAC domain, was divided into two intein-fused fragments that can be independently packaged into AAV vectors. Upon co-expression in neurons, intein-mediated splicing reconstitutes a full-length, functional biosensor.

As a proof of concept, we developed an AAV-compatible FRET biosensor for cAMP using a sequential design-and-validation workflow (**Fig. 1a**). Unlike previous approaches focused on delivering natural genes with large size [10,11], our work aims to deliver engineered biosensor integrating both biosensing functionality and optical performance. This workflow enables the conversion of conventional FRET-based biosensors into split, AAV-compatible designs by identifying structurally permissive split sites through domain analysis and protein structure prediction. Our work further incorporates fluorescent protein replacement to optimize optical performance, along with functional validation and reconstitution in cultured cells and mouse brains for diverse neuroscience applications.

**Figure 1.**
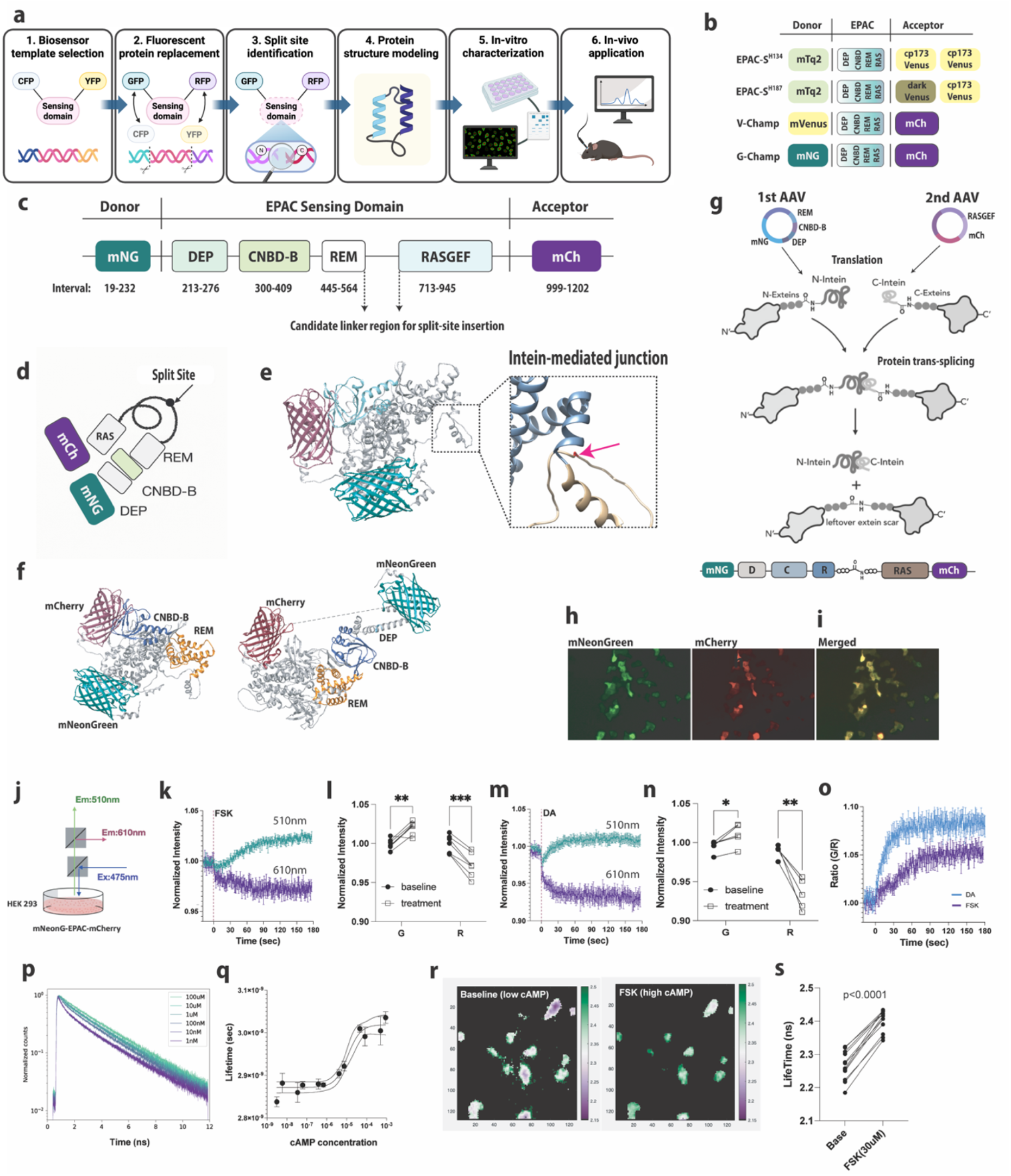
Split-protein reconstitution strategy for an AAV-compatible FRET cAMP biosensor and validation in HEK293 cells. (a) Overview of a sequential design and validation workflow for our cAMP biosensor. (b) Comparison between a conventional mTurquoise-Venus-based cAMP biosensor (EPAC-SH134, EPAC-SH187) and a redesigned mNeonGreen-mCherry cAMP-based biosensor optimized for one-photon in vivo applications. (c) Domain architecture of the complete biosensor, incorporating the EPAC sensing domain (DEP, CNBD-B, REM, and RasGEF) with mNeonGreen and mCherry. (d) Conceptual illustration of the linker region selected for intein-mediated splitting within the context of fluorescent proteins and EPAC subdomains. (e) AlphaFold-predicted structure supporting the idea that the split site lies within a noncritical linker region. (f) Predicted structures of complete biosensor in two different states, showing ligand-dependent conformational changes affecting donor–acceptor distance. (g) Two complementary biosensor plasmids are encoded on separate AAV-compatible constructs, which subsequently reconstitute inside the cell. (h,i) Fluorescence imaging of HEK293 cells following co-transfection with two plasmids. (j) Experimental schematic for intensity-based ratiometric characterization of the cAMP biosensor in HEK cells. (k) Forskolin (20 µM) induces time-dependent changes in green (510 nm) and red (610 nm) emission. (l) Two-way ANOVA of green (510 nm) and red (610 nm) intensities at baseline and post-treatment. Treatment effect: **p = 0.003; interaction: ***p = 0.0002; wavelength: p = 0.2318. Sidak’s test showed significant increases at 510 nm (**p = 0.0044) and 610 nm (***p = 0.002). (m,n) Dopamine (10 µM) stimulation in D1 receptor-expressing HEK cells similarly produces significant wavelength-dependent responses. Wavelength effect: *p = 0.0288; treatment effect: **p = 0.0022; wavelength × treatment interaction: ***p = 0.0003. Sidak’s multiple comparisons across treatment showed significant changes at 510 nm (*p = 0.042) and 610 nm (**p = 0.0064). (o) Green-to-red (G/R) emission ratio over time derived from these measurements. (p,q) Fluorescence lifetime measurements using 2-photon lysate characterization reveal cAMP-dependent increases in donor lifetime. (r,s) Fluorescence lifetime imaging in HEK293 cells shows a significant increase in lifetime following forskolin stimulation (two tailed paired t-test, p < 0.0001).

To implement this workflow, we selected an EPAC-based cAMP biosensor scaffold (**Fig. 1b**), as EPAC-SH134 [12]. A similar construct, EPAC-SH187, has been previously validated for in vivo imaging following utero electroporation [13]. The EPAC-based sensors used in these previous studies employed a cyan donor (CFP, mTurquoise2; extinction coefficient: 33,000 M^−1^cm^−1^; brightness: ∼28) and a yellow acceptor (YFP, Venus) for 2-photon imaging. However, we found that the CFP donor was suboptimal for one-photon fiber photometry or imaging due to relatively low photon counts. To improve excitation efficiency and compatibility with in vivo fiber photometry and one-photon imaging in freely behaving mice, we replaced CFP and YFP pairs with green and red excited fluorophores as the donor/acceptor pair (in this case mNeonGreen; extinction coefficient: 116,000 M^−1^cm^−1^; brightness: ∼93, and mCherry). The resulting mNeon**G**reen–m**Ch**erry-c**AMP** (**G-Champ**) biosensor incorporates EPAC1-derived domains (**Fig. 1c**).

Conserved domain analysis (see **Methods**) identified a non-critical linker (residues 565–712) between the REM and RasGEF domains of EPAC as a suitable split site (**Fig. 1c,d**). Structural modeling of full-length G-Champ further confirmed that this site lies within a flexible region and avoids structured domains (**Fig. 1e**). The predicted structure preserves REM and RasGEF architecture, consistent with the X-ray crystal structure [14], and provides a ∼30–40 Å change in donor-acceptor distance upon conformational transition (**Fig. 1f**). Based on this design, we divided the full-length biosensor into two fragments (mNeonGreen-EPAC-NT and mCherry-EPAC-CT), each encoded in separate AAV constructs (**Fig. 1f)**. We next validated biosensor reconstitution in HEK293 cells. Co-transfection of the two fragments (see **Methods**) was confirmed by co-expression of mNeonGreen and mCherry based on cell imaging (**Fig. 1h,i**).

We characterized biosensor function using intensity-based ratiometric fluorescence measurements (**Fig. 1j**, see **Methods**). Elevation of cAMP by forskolin (20 µM) or dopamine (10 µM, via D1R co-expression) produced robust changes in donor and acceptor signals, with faster kinetics observed for dopamine (**Fig. 1k–o**). With the same strategy, we could create the biosensor with mClover or mVenus as donors. These results demonstrate that **G-Champ** reliably reports cAMP dynamics in standard cell-based assays.

We next evaluated fluorescence lifetime responses of the **G-Champ** biosensor in cell lysates using a two-photon time-correlated single-photon counting (TCSPC) system (**Fig. 1p,q**, see **Methods**) by directly varying cAMP concentrations. In intact HEK cells, forskolin stimulation similarly produced measurable lifetime increases at the cellular level (**Fig. 1r,s**) using a fiber-bundle FLIM imaging system.

To assess in vivo performance, each biosensor fragment was packaged into separate AAV vectors and delivered to the mouse brain (**Fig. 2a**). Using transgenic mouse lines with Cre-dependent expression, we achieved cell type-specific targeting of expression to dopamine D1 or D2 receptor-expressing striatal medium spiny neurons (MSNs) as shown in **Fig. 2b-d**.

**Figure 2.**
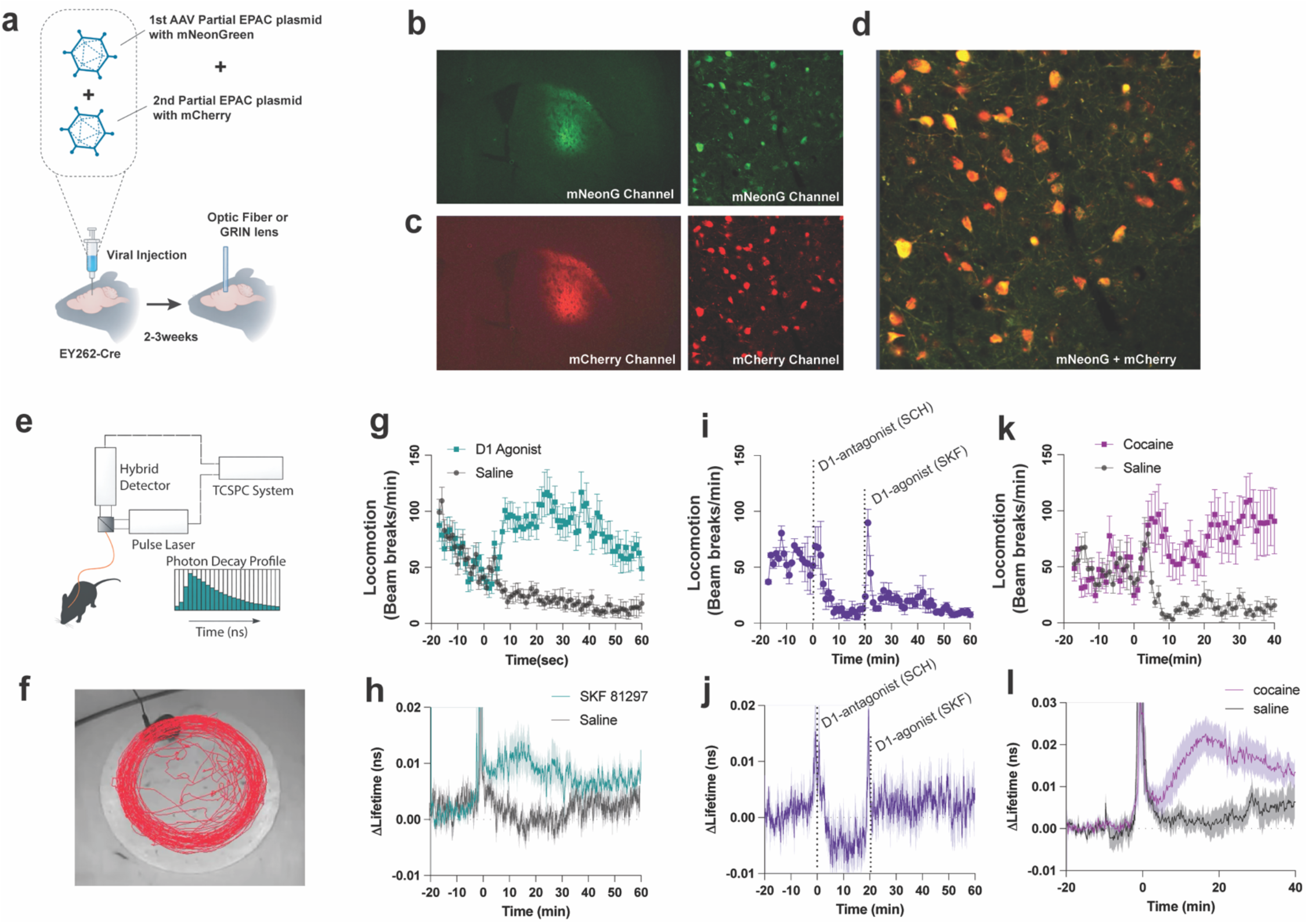
Viral delivery and in vivo reconstitution of cAMP biosensors for monitoring neural circuit signaling. (a) Dual-AAV strategy encoding complementary biosensor plasmids for in vivo biosensor reconstitution. (b-d) Representative fluorescence imaging of brain slices confirms Cre-dependent co-expression of the cAMP biosensor. (e) Diagram of the system used for in vivo fluorescence lifetime-based fiber photometry. (f) representative locomotor trajectories following administration of the D1 agonist SKF 81297 (7.5mg/kg). (g,h) SKF 81297 induces hyperlocomotion compared to saline controls and cAMP increase compared to the saline injection in the same subject. (i,j) Pharmacological validation shows that D1 receptor antagonist SCH 23390 (0.3mg/kg) blocks hyperlocomotion and agonist-induced cAMP elevation, confirming D1 receptor specificity. (k,l) Cocaine (20 mg/kg) produces robust increases in locomotion and cAMP levels in D1-MSNs.

We then evaluated functional responses in vivo using fluorescence lifetime-based fiber photometry in freely moving mice (**Fig. 2e**, see **Methods**). Intraperitoneal (I.P.) administration of the D1 receptor agonist SKF 81297 (7.5 mg/kg) induced a sustained increase in locomotor activity (**Fig. 2f,g**) accompanied by a corresponding elevation in cAMP levels in D1-MSNs (**Fig. 2h**).

To confirm receptor specificity, animals were pretreated with i.p. injection of the D1 receptor antagonist SCH 23390, which reduced both locomotor activity and cAMP signals and prevented further increases following agonist administration (**Fig. 2i,j**). Next, we examined the effect of i.p. injection of cocaine (20 mg/kg), which blocks dopamine reuptake and increases extracellular dopamine. Cocaine administration induced robust locomotor activity and an elevation of cAMP levels in D1-MSNs (**Fig. 2k,l**).

In summary, we present a split intein–based strategy that enables AAV delivery and in vivo reconstitution of oversized FRET biosensors. Applying this approach, we developed an EPAC-based cAMP sensor and demonstrated cell-type-specific monitoring of rapid and slow cAMP modulation in freely behaving mice. This platform facilitates the adaptation of FRET biosensors for in vivo use and expands access to quantitative, system-independent measurements of intracellular signaling.

## METHODS

### Domain analysis and protein structure prediction

Amino acid sequences of the biosensor constructs (provided in the Supplementary Information) were used for domain annotation and structural modeling. Conserved domain analysis was performed using the NCBI Conserved Domain Database (CDD; https://www.ncbi.nlm.nih.gov/Structure/cdd) to identify domain boundaries within the EPAC1-derived sensing region, including the DEP, CNBD-B, REM, and RasGEF domains.

Protein structure prediction was carried out using AlphaFold2 and AlphaFold3 (Google DeepMind; https://deepmind.google/science/alphafold/) on the NIH high-performance computing (HPC) system. Full-length biosensor constructs, including fluorescent protein fusions, were modeled to evaluate overall structural organization at residue-level resolution. The predicted models were analyzed to assess secondary structure elements, domain integrity, and the spatial positioning of candidate split sites relative to structured domain cores.

Structural visualization and analysis were performed using UCSF Chimera (https://www.cgl.ucsf.edu/chimera/) on a local workstation (Apple M2 Pro, 2023; 32 GB RAM; macOS Sequoia 15.5). The models were examined for overall folding, domain architecture, and the structural context of linker regions. Where applicable, predicted structures were qualitatively compared with available experimental structures (e.g., X-ray crystallography) to evaluate consistency in domain organization.

### Plasmid co-transfection in HEK293 cell

HEK293 cells were maintained in culture–treated flasks (BioLite™ Flasks, Thermo Fisher, #130189) under standard incubation conditions (37 °C, 5% CO2). Cells were cultured in DMEM (Dulbecco’s Modified Eagle Medium, Thermo Fisher, # 10569010) supplemented with 10% fetal bovine serum (Thermo Fisher, #A5209501) and 1% penicillin–streptomycin (Thermo Fisher, #15070063).

For experiments, cells were seeded into 12-well plates with 1 mL of complete DMEM. After 12 h, when cells reached approximately 50% confluency, transfection solutions were prepared by mixing 1 µg total DNA (mNeonGreen-EPAC-NT and mCherry-EPAC-CT plasmids) in 150 µL of DMEM (without fetal bovine serum or penicillin–streptomycin) with 4 µL of polyethylenimine (PEI). The culture medium in each well was replaced with 500 µL of fresh DMEM (4∼25 °C). The transfection mixture was then added to each well (50 µL per well). Plates were returned to standard incubation conditions (37 °C, 5% CO_2_) and maintained for 24–72 h prior to analysis.

### Ratiometric measurement of HEK293 cell

Culture medium (DMEM) was removed from the tissue culture flask, and cells were dissociated by adding 5 mL of TrypLE (Gibco, #12604-021), followed by incubation for 2–3 minutes at 25 °C in a laminar flow hood. An equal volume (5 mL) of fresh DMEM was then added, and cells were gently triturated to obtain a uniform cell suspension.

The resulting 10 mL cell suspension was transferred to a 15 mL Falcon tube and centrifuged at 800 × g for 4 minutes at 23 °C. After centrifugation, the supernatant was carefully aspirated, and the cell pellet was resuspended in 4–10 mL of fresh DMEM. Cells were seeded into 12-well plates and transfected with mNeonGreen-EPAC-NT and mCherry-EPAC-CT plasmids as described above. Once cells reached approximately 50% confluency, the culture medium was replaced with fresh DMEM. Cells were then detached by gentle pipetting, transferred, and seeded into clear-bottom, black-walled, tissue culture–treated 96-well plates. The plates were incubated at 37 °C in a humidified atmosphere containing 8% CO_2_ for 12–24 hours.

Before ratiometric characterization, medium was replaced with DPBS containing calcium and magnesium (Gibco, #14040-133), and baseline fluorescence (F_0_) was measured using a PHERAstar FSX microplate reader. cAMP elevation was induced with forskolin (20 µM) or dopamine (10 µM; with D1R co-expression). Changes in donor (510 nm) and acceptor (610 nm) fluorescence intensities were recorded for 180 s following drug application.

### 2-photon microscopy lysate characterization

The cell suspension was prepared from a tissue culture flask. Cells were then seeded into 12-well culture plates and co-transfected with mNeonGreen-EPAC-NT and mCherry-EPAC-CT plasmids as described above. After 3–5 days, co-expression of mNeonGreen and mCherry was confirmed by fluorescence microscopy (Zeiss Axiovert 200, with Zeiss FS 46 and FS 121)

Transfected cells were lifted from the 12-well plates using DPBS (Thermo Fisher, #14190144) supplemented with 10 mM EDTA and transferred to a 15 mL conical tube. Cells were centrifuged at 800 × g for 4 minutes at 23 °C. The supernatant was carefully removed, and the cell pellet was resuspended in 100 µL of lysis buffer (Promega Dual-Luciferase™ Reporter, Promega Corporation, #E194A) and 400 µL of deionized water (dH_2_O) for downstream analysis. The lysate was then centrifuged at 800 × g for 4 minutes to remove insoluble debris, and the clarified supernatant was collected for downstream analysis. The resulting lysate was diluted 1:10 in PBS and then mixed with varying concentrations of cAMP solution (Sigma-Aldrich, A6885) at a 1:1 ratio. For two-photon characterization, a wideband mode-locked Ti:Sapphire laser (Mai Tai, Spectra-Physics) was used at 950 nm. Fluorescence signals were detected using an HPM-100 Hybrid Detector Module and acquired with SPCM data acquisition software (Becker & Hickl).

### Fluorescence lifetime calculation

Fluorescence lifetime decay curves were fit using a biexponential model,

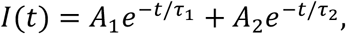

where *Ai* and *τi* denote the pre-exponential amplitude and fluorescence lifetime of the *i*-th component, respectively.

The intensity-weighted mean fluorescence lifetime (*⟨τ⟩*int) was calculated as:

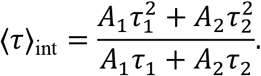

In our experimental results, A1,τ1,A2,τ2 are the primary parameters obtained from experimental TCSPC measurements of the cAMP biosensor at different cAMP concentrations.

### Animals

All animal procedures were carried out in compliance with the National Institutes of Health guidelines for the care and use of laboratory animals and were approved by the Animal Care and Use Committee of the National Institute on Alcohol Abuse and Alcoholism (NIAAA, protocol LIN-DL-1). Mice were maintained in the NIAAA animal facility under a 12-hour light/dark cycle with regulated temperature and humidity.

Surgical procedures, including viral injections and implantations, were performed on male and female mice aged 3 to 9 months. Wild-type C57BL/6J mice (Jackson Laboratory, stock #000664) were used for locomotor studies. Heterozygous Drd1-Cre mice [Tg(Drd1-cre)FK150Gsat/Mmucd] were obtained from GENSAT and bred with C57BL/6J mice. Adora2a-Cre BAC transgenic mice (GENSAT, KG139Gsat) and Chat-ires-Cre mice (B6.129S-Chattm1(cre)Lowl/MwarJ; Jackson laboratory, stock # 031661) were also crossed with C57BL/6J mice, and their offspring were used for the experiments.

Mice were housed either individually or in small groups (up to four mice per cage). All experiments were conducted during the light phase (06:00–18:00), and repeated measurements for individual animals were performed at consistent times across consecutive days.

### Virus injection and surgeries

To achieve Cre-dependent expression of the cAMP biosensor, two AAV vectors encoding mNeonGreen-EPAC-NT (titer: 6.9 × 10^12^ GC/mL) and mCherry-EPAC-CT (titer: 1.04 × 10^13^ GC/mL) were mixed at a 1:2 ratio and microinjected bilaterally (or unilaterally, where indicated) into transgenic mice. Stereotaxic coordinates were as follows: nucleus accumbens (NAc; AP +1.0 mm, ML ±0.7 mm, DV −4.7 mm from the brain surface) and dorsolateral striatum (DLS; AP +0.5 mm, ML ±2.2 mm, DV −3.2 mm from the brain surface).

For viral delivery, mice were anesthetized with isoflurane (1–2% in oxygen) and secured in a stereotaxic frame (Kopf Instruments). A midline scalp incision was made to expose the skull, and craniotomies were performed using a dental drill. Viral solution was injected using a 32-gauge Hamilton syringe (Hamilton, #65458-01) at a volume of 300 nL per site and a rate of 50 nL/min. Following injection, the needle was left in place for an additional 5–8 minutes to allow diffusion before slow withdrawal. The incision was then closed using VetBond tissue adhesive (3M, #1469SB). Mice were allowed to recover in heated cages for 3 days and then returned to their home cages for at least 2 weeks prior to secondary surgery.

For fiber implantation, mice were re-anesthetized and the implantable fiber optic cannula was slowly lowered into the target region through the existing craniotomy. The cannula (CFMC22L05, Thorlabs) was secured in place using a UV-curable adhesive (OptiBond−, Kerr Dental) and further stabilized with a dental cement headcap. Following surgery, mice recovered for at least 2 weeks, with ketoprofen administered for 3 days postoperatively, before subsequent experiments.

### In vivo fluorescence lifetime-based fiber photometry

At least three weeks after viral infusion, mice used for *in vivo* experiments underwent a second surgery for optical fiber implantation, following procedures similar to those previously described [15]. Fluorescence lifetime and intensity signals were sampled at 20 Hz during Pavlovian conditioning and pharmacology experiments, but at 3–7 Hz during long-term 24-hrs recording experiments. Laser excitation and emission collection were achieved using a custom-designed multimode patch cord (3m length, 200 µm core diameter, Thorlabs) with a numerical aperture (NA) of 0.22 and a 200 µm core diameter, terminating in a 2.5 mm ceramic ferrule. This ferrule was connected to an implanted fiber optic cannula (CFMC22L05, Thorlabs) via a ceramic mating sleeve.

Prior to the actual experiments, all animals were recorded for 5 minutes to assess signal quality, including absolute photon counts and whether the measured fluorescence lifetime reflected the biosensor signal rather than intrinsic tissue fluorescence. Animals with absolute photon counts lower than 1.5 × 10^5^ photons/s— comparable to intrinsic tissue fluorescence levels in the absence of fluorescent protein expression—were excluded from the animal experiments and analysis.

### Drug administration

SKF 81297 (D1 receptor agonist; Tocris, #1447) and SCH 23390 (D1 receptor antagonist; Tocris, #0925) were prepared from stock solutions in saline (vehicle) and administered via intraperitoneal (i.p.) injection. SKF 81297 stock solutions were gently warmed to facilitate dissolution. Cocaine hydrochloride (National Institute on Drug Abuse) was dissolved in sterile 0.9% saline on the day of the experiment and administered intraperitoneally at 20 mg/kg body weight.

## Acknowledgements

This research was supported by the Intramural Research Program of the National Institutes of Health (NIH). The contributions of the NIH authors are considered Works of the United States Government. The findings and conclusions presented in this paper are those of the authors and do not necessarily reflect the views of the NIH or the U.S. Department of Health and Human Services.

## Notes

### Competing Interest Statement

The authors have declared no competing interest.

